# Single Cell Transcriptomics Reveals Global Markers of Transcriptional Diversity Across Different Forms of Cellular Senescence

**DOI:** 10.1101/2021.06.16.448710

**Authors:** Shane A. Evans, Yee Voan Teo, Kelly Clark, Takahiro Ito, John M. Sedivy, Nicola Neretti

## Abstract

Cellular Senescence is a state of irreversible cell cycle arrest, and the accumulation of senescent cells contributes to age-related organismal decline. The detrimental effects of cellular senescence are due to the senescence associated secretory phenotype (SASP), an array of signaling molecules and growth factors secreted by senescent cells that contribute to the sterile inflammation associated with aging tissues. Recent studies, both in vivo and in vitro, have highlighted the heterogeneous nature of the senescence phenotype. In particular, single cell transcriptomics has revealed that Oncogene Induced Senescence (OIS) is characterized by the presence of subpopulations of cells expressing different SASP profiles. We have generated a comprehensive dataset via single-cell transcriptional profiling of genetically homogenous clonal cell lines from different forms of senescence, including OIS, Replicative Senescence (RS), and DNA Damage Induced Senescence (DDIS). We identified subpopulations of cells that are common to all three major forms of senescence and show that the expression profiles of these subpopulations are driven by markers formerly identified in individual forms of senescence. These common signatures are characterized by chromatin modifiers, inflammation, extracellular matrix remodeling, and Ribosomal protein expression. The expression patterns of these subpopulations recapitulate primary and secondary senescence, a phenomenon where a preexisting (primary) senescent cell induces senescence in a neighboring (secondary) cell through cell-to-cell contact. Since it is still unclear what type of senescence occurs in-vivo with age, it is important to know that the formation of primary and secondary populations is common to multiple types of senescence since this mechanism could help explain how senescent cells accumulate in aged organisms. Finally, we show that these subpopulations show differential susceptibility to the senolytic agent Navitoclax, suggesting that senolytic agents targeting the apoptotic pathways may be clearing only a subset of senescent cells based on their inflammatory profiles in-vivo.

## Introduction

Cellular senescence (CS) is a programmed stress response that leads to a cell’s permanent exit from the cell cycle and can be induced by a variety of factors including telomere attrition, oncogene activation, oxidative stress, and DNA damaging agents [1–3]. Although CS comes in different forms, an established senescent pathway involves a persistent DNA damage response that leads to the activation of the tumor suppressor protein 53 (P53), which in turn activates the cyclin dependent kinase inhibitor gene CDKN1A. The translated protein encoded by the CDKN1A gene, p21, holds the cell in cell cycle arrest until upregulation of the cyclin dependent kinase inhibitor gene CDKN2A, which encodes the p16 protein that maintains the cell in an irreversible senescent state [1, 4]. During senescence, the cell undergoes global epigenetic changes including dramatic chromatin alterations and increased expression of an array of extracellular remodeling proteins, growth factors and inflammatory molecules such as interleukins and interferons that compose the senescence associated secretory phenotype (SASP) [4–7]. SASP leads to inflammation disrupting the tissue microenvironment, reinforces the senescent phenotype by contributing to the cell cycle arrest, and can induce paracrine senescence in normal cells [8, 9].

Studies describing the heterogeneity of senescent cells have shown that there are different forms of SASP. For example, one form is dominated by Transforming Growth Factor Beta (TGF-beta) signaling [10–12]. TGF-beta dominated profiles are characterized by extra-cellular matrix remodeling, collagen deposition, and by the expression of growth factors such as the Connective Tissue Growth Factor (CTGF) [10, 13, 14]. Another form of SASP however, shows a more proinflammatory profile with higher levels of interleukins such as IL1A, IL1B, IL6 and other genes regulated by NFKB [10–12, 15]. In OIS, the prevalence of these profiles changes as cells persist in the senescent state: TGF-beta signaling is typically higher in the stages of senescence, while in the later stages senescent cells transition to a pro-inflammatory phenotype [10].

Not only do transcriptional profiles change over the course of time, but they can also characterize different types of senescent states. For instance, TGF-beta dominated SASP profiles are prevalent in Notch Induced Senescence (NIS) [10–12], which occurs when a primary senescent cell makes direct contact and activates the Notch signaling pathway in a neighbor cell, causing the spread of senescence through juxtacrine signaling to secondary senescent cells. OIS cells can act as the primary senescence source, and display more pro-inflammatory SASP profiles with higher levels of interleukins and NFKB regulated genes in contrasts with the anti-inflammatory TGF-beta dominated profiles of NIS cells [11].

Cellular senescence is an inherently heterogeneous state, as it can be cell type- and insult-dependent [1, 4, 16]. However, most of the data describing senescence has been collected using bulk sequencing technologies that measure average gene expression across large heterogeneous pools of cells and is blind to cell-to-cell transcriptional variability. With the advent of single cell transcriptomics, it has become clear that distinct subpopulations contribute to the population-wide average [11, 17]. Despite the limited single cell transcriptional data currently available for senescent cells, it is becoming increasingly clear that senescent cells are subject to significant transcriptional diversity [10, 16–18]. Consistent with this transcriptional heterogeneity, subpopulations of senescent cells have been observed in single cell transcriptomic studies from fibroblasts and endothelial cell lines [11, 17, 18]. For example, using single cell RNA-seq, Teo et al. showed that two subpopulations co-exist in the OIS cultures [11]. One population possessed the familiar OIS transcriptional profile (primary senescence), while the second population, which was also senescent, possessed SASP profiles that were growth factor rich and dominated by TGF-beta signaling, and was composed of NIS cells (secondary senescence).

An important difference between OIS and NIS is the expression of HMGA1 [12, 19, 20]. HMGA1 is a highly abundant chromatin associated protein that is an essential component of the senescent chromatin architecture and is critical for the onset of the senescence program in OIS. OIS typically upregulates HMGA1, and this creates dramatically different chromatin structures compared to NIS cells [12]. Even though HMGA1 is a very important protein for the execution of the senescence program and the organization of the senescent chromatin, HMGA1’s role in senescence and its opposition of anti-inflammatory and TGF-beta dominated SASP profiles has almost entirety been explored in the context of OIS and NIS.

The senescent phenotype is characterized by chromatin modifications, DNA-damage response pathways, and SASP profiles [1, 3]. However, an in-depth description of how these different aspects vary across senescent cell populations is still lacking. Here, we used single cell transcriptomics [17, 21] to identify subpopulation of senescent cells in genetically homogenous clonal cell lines with a inducible HRAS:G12V transgene that was activated only in OIS cultures. This strategy ensures that the subpopulations we observe are not a caused by variability of the number and genomic location of the transgene constructs, and the consequent variability in the expression levels of the transgene across cells. This experimental design allowed us to conduct a novel, unified analysis of senescent cell heterogeneity across the major forms of senescence.

## Results

### Single-cell Transcriptional Profiling of Clonal Lines

We performed single cell RNA-sequencing using the 10x Chromium microfluidics platform to study cell-to-cell gene expression heterogeneity in RS, OIS, and DNA Damage Induced Senescence (DDIS). All cell populations were generated using female human diploid fibroblast cells (LF1 cells) and were derived from a clonal cell line that possessed a 4- Hydroxytamoxifen (4-OHT) inducible HRAS:G12V transgene (Figure 1A). Although subpopulations of senescent cells had been identified in OIS cultures, little is known about the cell-to-cell diversity in other forms of senescence. We hypothesized that the diversity found in OIS cultures is a phenomenon common across multiple forms of senescence. To test this hypothesis we sequenced different types of senescence in different cell culture conditions.

**Figure 1.**
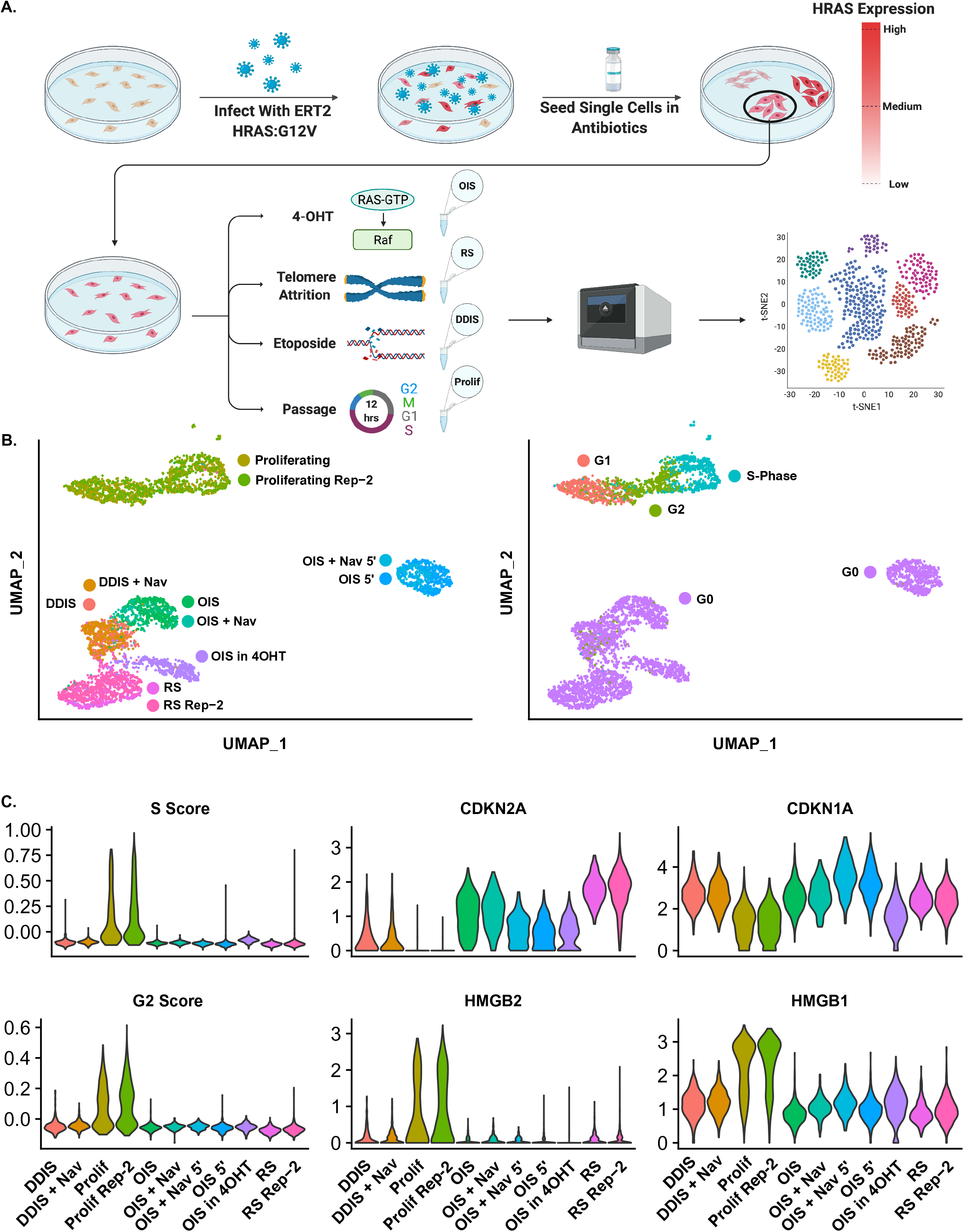
Experimental Overview. **A.** Schematic showing generation of clonal cell lines with HRAS:G12V transgene and overall experimental design. **B.** UMAP plots showing all data sets generated for this study (left) and their predicted stage in the cell cycle (right). **C.** Violin plots showing down regulation of cell cycle genes, HMGB1, HMGB2 and upregulation of CDKN2A and CDKN1A in senescent cultures.

For OIS cultures we have libraries that were in 4-OHT, while other were removed from 4-OHT prior to sample collection. This helped us to determine how 4-OHT affected diversity. For RS we sequenced two clonal cells lines, referred to as RS and RS Rep2. This let us show that the diversity we saw in our datasets was not due to the effects of a single clonal cell line but instead could be recapitulated across multiple clonal lines. We also have DDIS and OIS cells that were treated with Navitoclax prior to sequencing. These datasets showed how the subpopulations we identified responded to senolytic drug treatment. Moreover, for OIS samples we also performed 5’ end sequencing. This let us show that the diversity we discovered in our datasets is not due to a 3’ sequencing bias. For full list of culture conditions sequenced for this publication see **Figure 1 B**. For the main figures in this publication we focus on the three of the data sets comprising RS, OIS, and DDIS cells. Analysis of the remaining data sets, which revealed strikingly similar diversity, are shown in the supplemental figures.

Since all of our datasets were generated with cells that were infected with a HRAS:G12V transgene, controlling for variable expression of this transgene is important if we are to draw conclusions regarding the formation of subpopulations in senescence. Without clonal cell lines the transgene would be expressed at varying levels across the cells, and since HRAS is upstream of many important molecular pathways that are heavily implicated in senescence, then this would have been a confounding factor in our experiments. This methodology contrast previous studies which conducted scRNA-seq on non-clonal cell lines that variably express the transgene [11]. Moreover, working with clonal cell lines ensured that any variability in gene expression that we observed was a product of the senescent phenotype and not due to a pre-existing heterogeneity in the proliferating cell populations.

For all types of senescence we noticed the presence of a large populations of cells that showed signs of low-quality data. These included higher than average expression of Ribosomal-RNAs in combination with low number of genes being expressed, and low number of unique molecular identifiers (UMIs). Additionally, we saw a different and smaller subpopulation of cells with high number of mitochondrial reads. Our proliferating control cells did not possess these subpopulations in large numbers. Taken together, these observations suggest that senescent cells are more fragile to the microfluidics used to generate these libraries. For all dataset we filtered out these low-quality data points and the remainder of our analysis focused on the higher-quality cells which we retained. See methods section and **Supplemental Figure 7** for a detailed explanation of our filtering strategy. RS, OIS, and DDIS cultures showed clear senescent gene expression patterns including upregulation of CDKN2A and downregulation of cell cycle genes, HMGB1, and HMGB2 (**Figure 1 C, Figure 2 H-J**).

**Figure 2.**
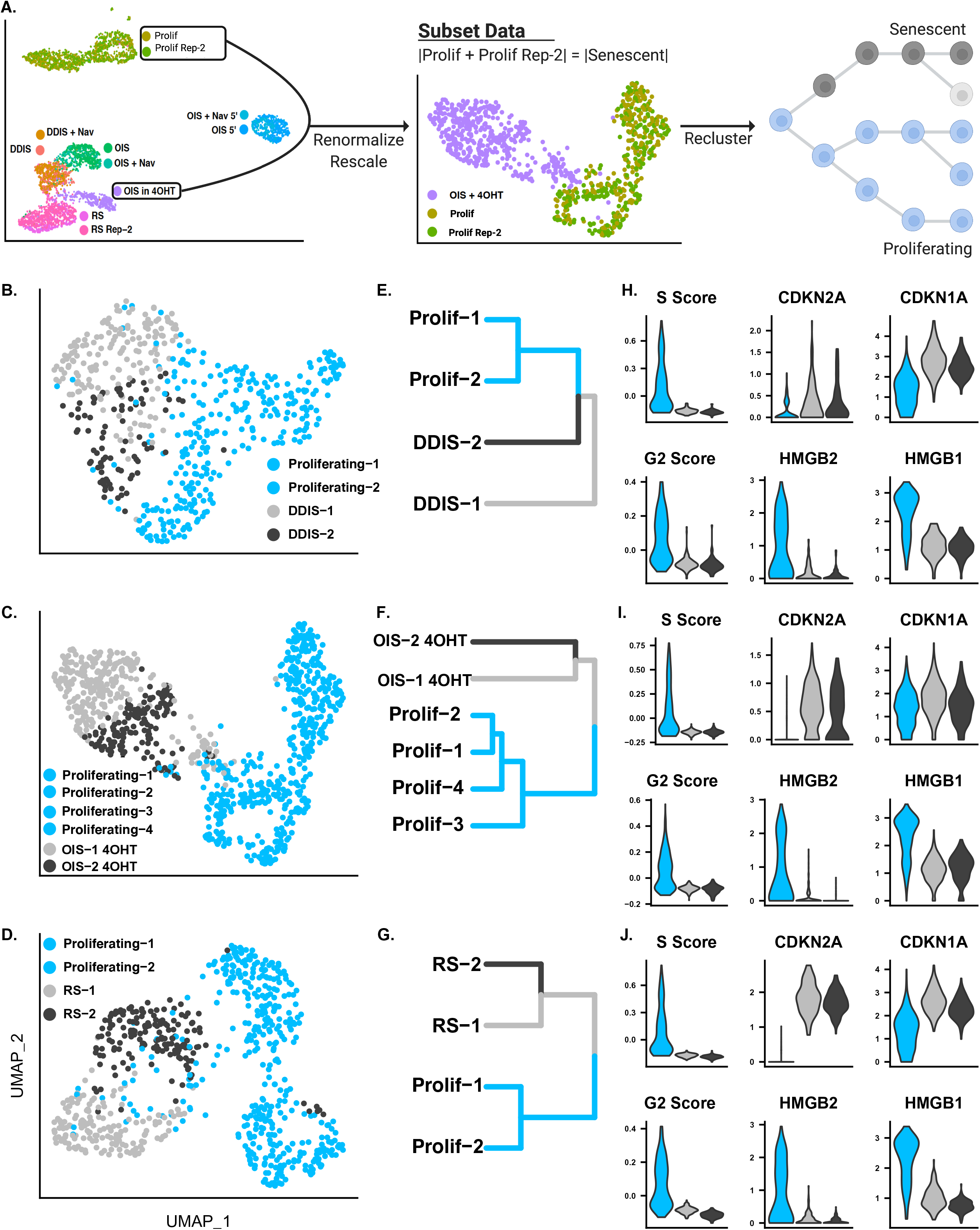
Senescent Cells Are Composed of 2 Clusters. **A.** Schematic showing the process for analyzing each senescent data set seperatley. OIS data was chosen for this illustration, but the same process was used for all data sets. **B. - D.** UMAP plots showing DDIS, OIS, and RS clusters. **E. - G.** The corresponding cluster trees. **H. - J.** Violin Plots showing that similar levels of senescence markers for each subpopulation.

All data sets were merged together using the Seurat Bioconductor Package [21, 22]. Our filtered data set included a total of 6,108 cells split across multiple types of senescence and conditions. We projected individual cells’ transcriptional profiles onto two dimensions using the Uniform Manifold Approximation and Projection reduction (UMAP) and observed a clear separation between senescent and growing cells (**Figure 1 B**). Computing a cell cycle score with Seurat shows that the proliferating population are composed of cells in the S, G2/Mitosis, and G1/G0 phases of the cell cycle. We noticed the different types of senescent cells are positioned more closely to control cells in the G1/G0 phase, showing that the senescent state as a distinct and final cell cycle phase (**Figure 1 B**).

### Clustering Reveals Two Subpopulations in Clonal Cell Lines

We combined each senescent data set along with an equal sized subpopulation of control cells (**Figure 2 A**). Clustering was performed on the resulting data sets (see **Methods**), and in all cases, we could identify subpopulations of senescent cells. For each type of senescence we call the subpopulations ‘-1’ and ‘-2’ (**Figure 2 B-G.**). We analyzed the expression of the transgene between the two clusters by aligning to the amino 3ʹ-glycosyl phosphotransferase (neo) selectable marker gene. We saw that the transgene is much more evenly expressed between the subpopulations we observe in cultures generated from clonal cell lines when compared to previous published experiments which did not use clonal lines (**Supplemental Figure 6**).

### Subpopulations are Distinguished by TGF-beta Signaling, DNA Damage Response, and Inflammatory Pathways

We performed a differential expression analysis between Cluster-1 and Cluster-2 in each type of senescence using the Seurat package with MAST methodology [23, 24]. The computed log fold changes of genes passing a false discovery rate of 0.05 were used to identify potential upstream regulators via the Ingenuity Pathway Analysis (IPA) (**Figure 3 C**). We plotted the upstream regulators according to two dimensions. On the x-axis are the computed z-scores, which show the direction of regulation. On the y-axis is the negative logarithmic transformation of their false discovery rate, which determines the upstream regulators whose gene sets show a statistically significant overlap with the list of differentially expressed genes in the data.

**Figure 3.**
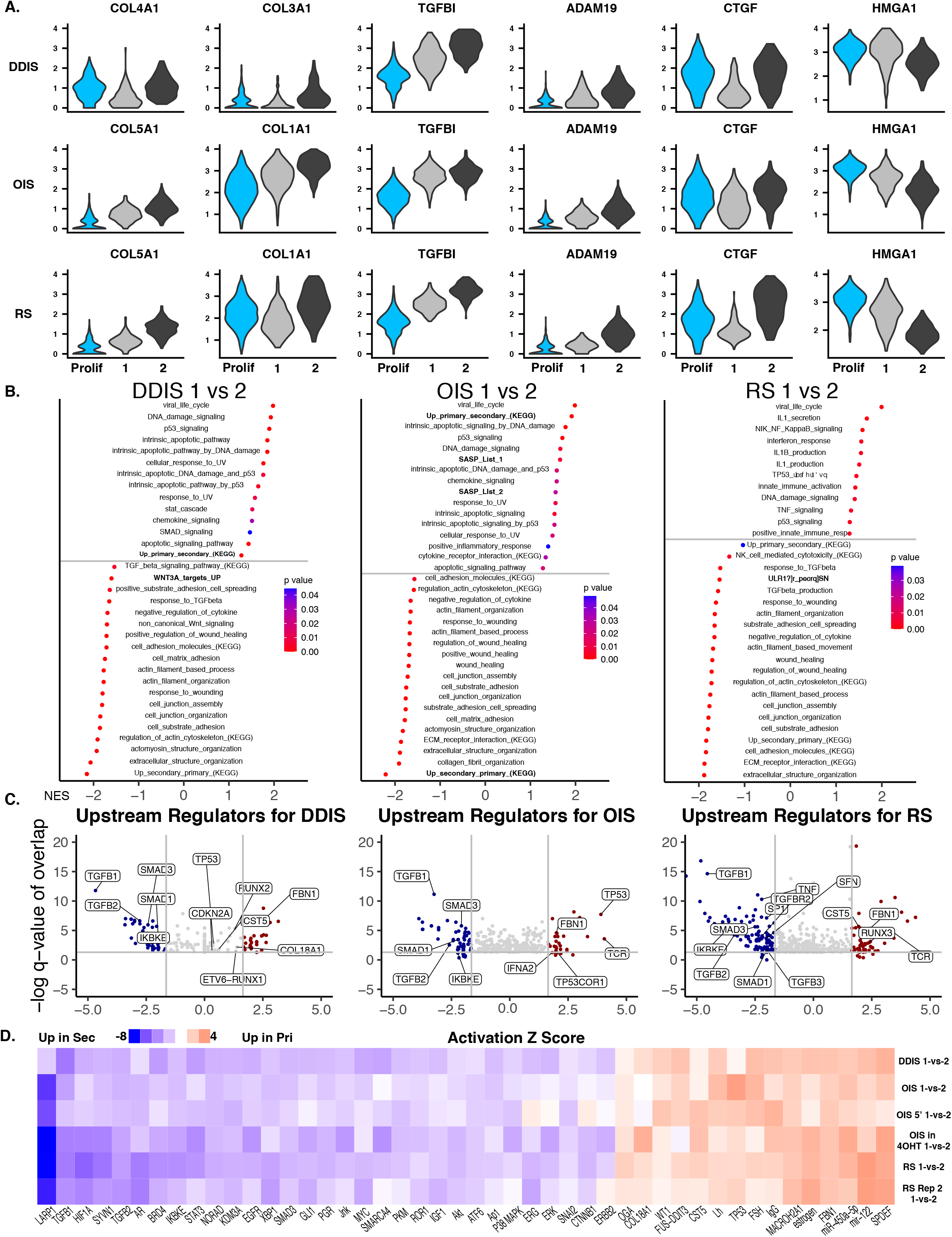
Cluster-1 and Cluster-2 Reveal Features of Primary and Secondary Senescence Across All Froms of Senescence. **A.** Violin plots showing expression of markers genes for primary and secondary senescence for cluster-1 and cluster-2 in DDIS, OIS, and RS cells. **B.** KEGG and GO terms (KEGG and GO analyses were run seperatley) plotted according to their NES scores along the x-axis and colored according to the p-value. **C.** Voclano pots showing predicted upstream regulators (IPA) for cluster-1 (red) and cluster-2 (blue) for each type of senescence. Regulators are plotted according to their z-score on the x-axis which shows if they regulate cluster-1 or cluster-2. The ‘-log of the q-value of overlap’ is plotted on the y-axis. This is the negative logarithmic transformation of their false discovery rate, which determines the upstream regulators whose gene sets show a statistically significant overlap with the list of differentially expressed genes in the data.

Using a z-score cut off 1.64 and –log q-value cut off of 1.301, we were able to identify statistically significant upstream regulators for each type of senescence. We noticed that the predicted upstream regulators were very similar for each type of senescence. More specifically, Cluster-1 showed upstream regulators TP53, a marker for DNA-damage [13, 25–27]. We also saw Interferon term IFNA2, suggesting a more proinflammatory profile [28]. Meanwhile, Cluster-2 showed upstream regulators corresponding to TGF-beta signaling and extra-cellular matrix remodeling. These regulators included TGF-Beta receptors, TGF-Beta1-3, and several SMAD proteins that are transducers for TGF-Beta signaling. We also performed a comparison analysis for each type of senescence and noticed a strong similarity in the z-scores for the upstream regulators (**Figure 3 D**) between all types of senescence that we sequenced. We also confirmed the existence of these subpopulations in all other senescent data sets (**Supplemental Figure 1** and **Supplemental Figure 9**).

Next, we took the log fold changes between cluster-1 and cluster-2 for each type of senescence and conducted a pre-ranked Gene Set Enrichment Analysis (GSEA) with Gene Ontology (GO) terms and Kyoto Encyclopedia of Genes and Genomes (KEGG) pathways [29–34]. We plotted the Normalized Enrichment Scores (NES) for statistically significant (p<0.05) GO terms and KEGG pathways. We noticed that subpopulations in Cluster-1 showed enrichment for terms related to DNA damage response pathways such as Telomere Organization, Mismatch Repair, Base Excision Repair, and DNA-ligation. Moreover, there was also an enrichment for SASP and inflammation pathways including our Custom SASP lists (See **Methods** for details on custom SASP lists), inflammatory response, cytokine interactions, viral life cycle, neutrophil migration, and viral transcription (**Figure 3 B**).

In contrast, cells in Cluster-2 showed higher levels of extra cellular matrix activity including terms related to integrin signaling pathways, extracellular organization, cell adhesion, collagen organization, MHC class 1 antigen presentation, and cell junction organization. Cluster-2 was also enriched for terms related to TGF-beta signaling such as regulation of TGF-beta, SMAD protein signaling, WNT signaling and anti-inflammatory gene activity (**Figure 3 B**). Collagen deposition, extra cellular matrix remodeling, and WNT signaling are all regulated or co-activated by TGF-beta signaling [13, 14, 25, 35]. Therefore, this is consistent with the expression profile of cluster-2 cells as being dominated by TGF-beta signaling pathways. These GO terms and KEGG pathways were also verified in all other data sets we collected (**Supplemental Figure 8**).

### Subpopulations Differentially Express HMGA1 and Resemble Primary and Secondary Senescent Cells

We investigated whether there were any differences in the expression levels of important chromatin modifiers. We found that HMGA1 is consistently upregulated in Cluster 1 (**Figure 3 A**). The HMGA1 gene encodes a highly abundant chromatin associated protein that has been shown to organize chromatin architecture in senescence and is critical for the onset of OIS. Moreover, previous studies have shown that HMGA1 is expressed higher in OIS when compared to NIS, which is characterized by increased levels of TGF-Beta signaling. This difference in HMGA1 expression was found to be responsible for many of the differences in chromatin architecture that exist between OIS and NIS cells, and many of these differences are predictive of gene expression [10–12]. These studies comparing NIS and OIS, along with the fact that we see TGF-Beta signaling and elevated HMGA1 expression arising from two distinct subpopulations in our scRNA-seq data, suggests that these HMGA1 expression is elevated only in the senescent cells where TGF-Beta expression is low and levels of DNA damage are high.

Since HMGA1 is differentially expressed between OIS and NIS cells, we decided to compare our data to the single cell transcriptional profiles from Teo et al.[11], as they showed that OIS cultures generated from non-clonal cell lines are composed of primary and secondary senescent subpopulations. For all four data sets (RS, OIS, DDIS, and Teo et al.) we see differential expression of collagen genes, TGF-beta response genes, and HMGA1 which are markers for primary and secondary senescence (**Figure 3 A** and **Supplemental Figure 4**). Moreover, the expression of HMGA1 is anti-correlated with TGF-beta signaling. We then ran a KEGG analysis with terms generated by Teo et al. relating to primary and secondary senescent phenotype, we see that our subpopulations are enriched for these terms, referred to as “UP_primary” and “UP_secondary” respectively. Like with our own data, we conducted a GSEA analysis for the data generated by Teo et al. and identified very similar GO terms and KEGG pathways. Cluster-1 is enriched for SASP, Inflammation, and DNA-damage response pathways. Meanwhile, Cluster-2 is enriched for extra-cellular matrix remodeling, Collagen deposition, and WNT signaling (**Supplemental Figure 8**).

### Subpopulations Show Differential Sensitivity to the Senolytic Agent Navitoclax

We wanted to determine if we could characterize the subpopulation of cells that are more resistant to treatment with BH3 mimetics. We treated DDIS and OIS cells with 1uM of the BH3 mimetic Navitoclax for three days [36]. Cells were then harvested and run on the 10x chromium platform to generate single cell libraries along with DMSO treated controls, see methods and **Figure 4 A** for Navitoclax experimental schematics. Data was aligned and filtered as previously described.

**Figure 4.**
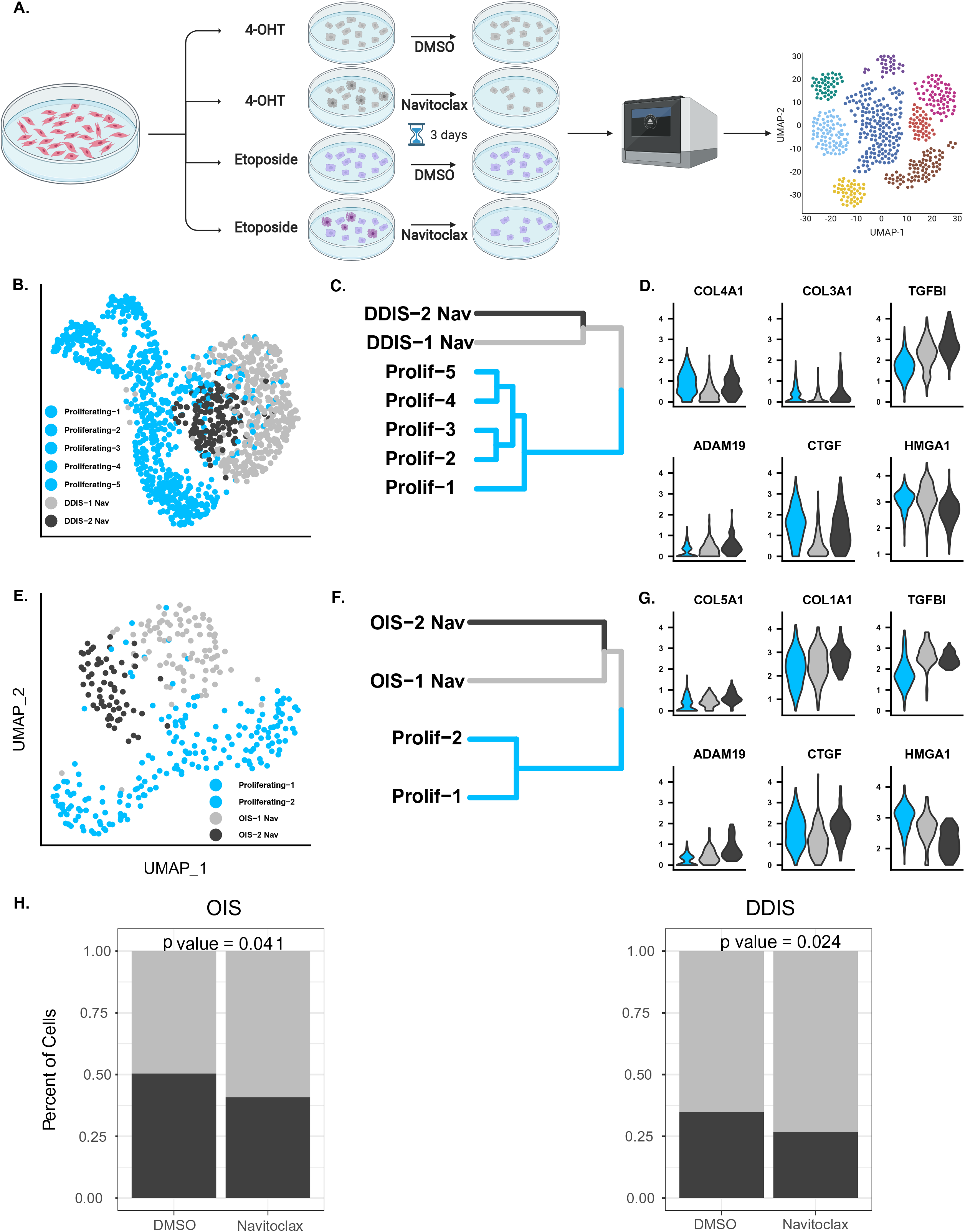
Navitoclax Preferentially Kills Cells in Cluster-2. **A.** Schematic showing experimental design for the Navitoclax experiments conducted on OIS and DDIS cells. **B. - D.** UMAP plots, cluster trees, and violin plots of primary and secondary senescence for DDIS cells treated with Navitoclax. **E. - G.** UMAP plots, cluster trees, and violin plots of primary and secondary senescence for OIS cells treated with Navitoclax. **H.** Bar plots showing that Navitoclax preferentially induces apoptosis in cluster-2 cells for OIS and DDIS cells.

We identified the cluster-1 and cluster-2 in Navitoclax treated samples as well (**Figure 4 B-G**). We observed that Navitoclax preferentially induced apoptosis in DDIS and OIS cells in Cluster-2, the subpopulation enriched for a secondary senescence, TGF-beta signaling, extracellular matrix remodeling, and lower levels of DNA-damage. (**Figure 5 A-B**). Moreover, we also saw this phenomenon in OIS cells that were sequenced from the 5’ end, although this data set did reach statistical significance (**Supplemental Figure 3**). This suggests that TGF-beta signaling in senescence may sensitize cells to drug induced apoptosis, emphasizing the translational importance of these subpopulations. In general these data demonstrate that for the development on senolytic agents it is important to consider the inflammatory profile of the target cell.

**Figure 5.**
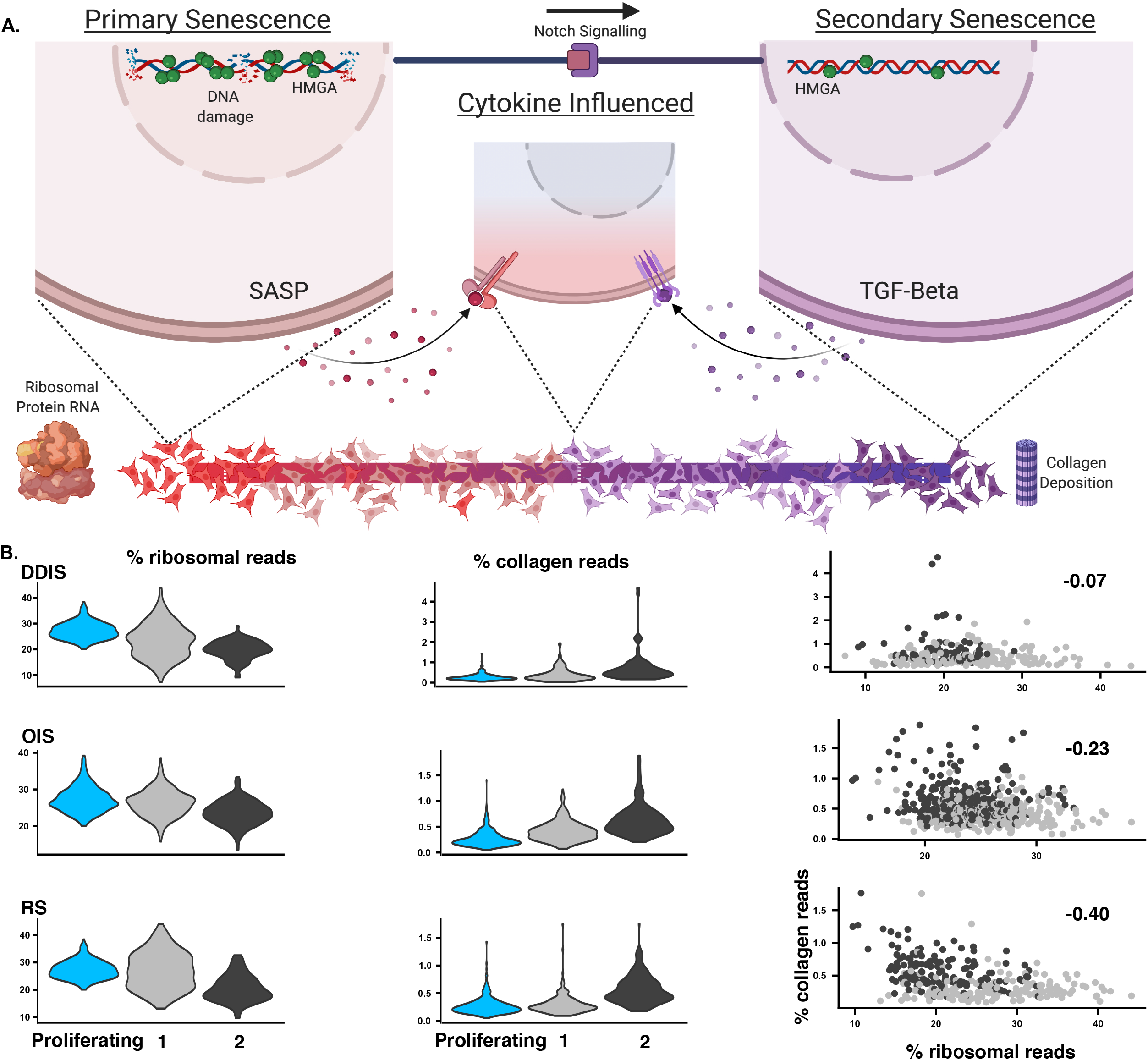
Cluster-1 cells Accumulate Ribosomal Transcripts and Cluster-2 Expresses Collagen Genes. **A.** Model showing explaining the observed expression profiles in our data sets. **B.** (Left) Violin plots showing the expression of collagen and ribosomal genes in cluster-1 and cluster -2 for RS, OIS, and DDIS cells. (Right) scatter plot showing that cells expressing higher levels of ribosomal reads express lower levels of collagen reads. Cells are colored according to their cluster-1 and cluster-2 assignments.

### Subpopulations Differentially Express Collagen and Ribosomal Protein Genes

Another feature we noticed in our analysis is that cells belonging to cluster-1 significantly accumulate ribosomal transcripts. Altered ribosome biogenesis is implicated heavily in senescence [37], and we extend these findings further by showing that it is a process differentially regulated between subpopulations. We plotted cells according to their expression of ribosomal genes on the x-axis and expression of collagen genes on the y-axis and saw a strong anticorrelation for all data sets, suggesting that cluster-2 and secondary senescent cells are characterized by the high levels of collagen genes and cluster-1 cells express high levels of ribosomal protein genes (**Figure 5**).

### Subpopulations Expression Profiles are Identified in Senescent Endothelial Cells

Since we identified these subpopulation in multiple forms on senescence in our clonal cell lines in addition to the IMR90 experiments conducted by Teo et al, we wanted to verify if these diversity profiles existed in another cell type. We downloaded single cell transcriptomic data sets generated from HUVEC cell lines [18]. These data were collected from cells as they transitioned into RS. This means that there were many more subpopulations in the HUVEC data set and it was difficult to distinguish senescent form pre-senescent cells. However we were still able to show major features of the diversity we saw in our own data.

We wanted to see if HUVEC cells expressing higher level of HMGA1 expressed low levels of collagen. We plotted cells according to their expression of HMGA1 (markers for cluster-1 and primary senescence) on the x-axis and expression of collagen genes (marker of cluster-2 and secondary senescence) on the y-axis. We saw an anticorrelation between these two features. We also plotted cells according to their expression of ribosomal reads (which we saw higher in cluster-1) on the x axis and saw a strong anticorrelation with cells that expressed high level of collagen reads. Moreover, reads mapping to ribosomal genes also anticorrelate with cells that have high number of reads mapping to collagen genes. See **Supplemental Figure 5** for this analysis.

## Discussion

Our analysis compares subpopulations of senescent cells across different forms of senescence. We observe that DDIS, RS, and OIS cells are composed of two subpopulations. For our experiments, we used clonal cell lines that were infected with a 4-Hydroxytamoxifen HRAS:G12V inducible transgene that was only activated in OIS cultures. The fact that we have discovered subpopulations of senescent cells being generated from clonal cell lines suggests that the formation of these subpopulations is an inherent property of the senescent phenotype, and not due to a preexisting heterogeneity that was present in proliferating cells. Therefore, these results offer additional insight into the formation of senescent cell subpopulations when compared with pre-existing scRNA-seq studies that have been generated from non-clonal cell lines. For instance, scRNA-seq studies on OIS samples that were infected with an inducible HRAS construct and then selected for with antibiotics yields a population of cells with varying amounts of HRAS expression, that ends up influencing the diversity of the final senescent population.

It is important to mention that due to the experimental conditions, our cell yield in the various forms of senescence was different, which affected the sensitivity with which we could detect differential expression between the two clusters. While this did not compromise our ability to detect common signatures, some of the genes that appear to be specific to one form of senescence and not the others might be the result of false negatives due to low cell numbers and statistical power.

For all forms of senescence, we have identified a subpopulation of cells that shows higher levels of extracellular matrix remodeling genes and genes that are regulated by TGF-beta such as collagen genes. Meanwhile, the other subpopulation shows higher levels of DNA-damage, pro-inflammatory profiles, and HMGA1 expression, and a strong accumulation of ribosomal transcripts. These molecular signatures are reminiscent of what is seen in senescent cells inducing Notch senescence in neighboring cells [10–12]. Consistently with this model, we see that the subpopulations we observe show similar gene expression patterns as described in the study conducted by Teo et al. [11] where a primary population of cells was driven into the senescent state through the activation of the HRAS oncogene, and in turn drove neighboring cells into senescence through the Notch signaling pathway (juxtacrine-induced senescence), giving rise to the second cluster of cells characterized by the activation of the TGF-beta signaling pathway [11]. Our study extends this work by showing that these subpopulations appear to be a universal feature of senescence as they consistently appear in clonal cell lines and in all the main forms of senescence. One possible explanation is that that ability for OIS cells to induce secondary senescence though the Notch signaling pathway is also shared by RS and DDIS cells (**Figure 5 A**).

Another possibility explaining the formation of these subpopulations is that that senescent cells tend to express TGF-beta early on in their senescent lifespan, while expressing higher levels of inflammatory and interferon response genes later in their lifespan [10]. Our analysis could be capturing a transition from a senescence phase dominated by TGF-beta signaling to a phase displaying a more pro-inflammatory profile. If this is the case then HMGA1 offers an interesting candidate whose role in this transition could be further investigated. As mentioned previously, HMGA1 has been shown to be downregulated in secondary senescent cells induced through a Notch signaling pathway when compared to their primary OIS counterparts. NIS is also characterized by high levels of TGF-beta signaling, which is consistent with a model where HMGA1 is implicated in the expression of TGF-beta pathways. Another interesting possibility is that the heterogeneity we are seeing is a result of differences in levels of DNA-damage [10–12, 19]. We noticed that cells in Cluster-1 are enriched in pathways relating to the DNA damage response. Therefore, higher levels of DNA-damage could predispose a cell to displaying the molecular signatures that we see in Cluster-1 such as increased SASP, lower TGF-beta signaling, and higher levels of HMGA1. Therefore, our study sets the stage for important questions. For instance, how does HMGA1 interact with DNA-damage response pathways in cellular senescence?

We also see that these subpopulations are translationally important by showing that treatment with the commonly used senolytic agent, Navitoclax, preferentially kills cells in Cluster-2. A possible explanation for this is that the higher levels of TGF-beta signaling, which is a known affecter of the apoptotic pathways [38], sensitizes cells to the apoptotic effects of Navitoclax [36]. Another explanation is that cells in Cluster-1 are at a later stage of senescence, and therefore are more resistant to apoptosis and senolytic drugs. It is also possible that cells in Cluster-1 are indeed primary senescent cells, and Navitoclax resistance is an inherent property of primary senescence. In any of these cases, this study points to the translational importance of senescent cell heterogeneity.

## Methods

### Single Cell RNA-sequencing

Human diploid fibroblast cells were trypsinized and centrifuged at 500 rcf for 10 minutes. Cells were re-suspended in cold Phosphate Buffered Saline and passed through a 40uM Flowmi Cell Strainer. Cells were then counted and loaded onto the 10x Chromium using V2 chemistry of 10x Genomics’ 3-prime Single Cell Reagents. Libraries were prepared according to the manufacturer protocol, and sequenced on a Hi-Seq platform at GeneWiz with manufacture recommend sequencing specifications. Senescent libraries were subjected to two rounds of sequencing. All single cell RNA-seq datasets generated in this work have been deposited in the Gene Expression Omnibus (GEO) database with accession number GEO:XYZ (Pending).

### scRNA-seq Data Processing and Filtering

Cell specific barcodes were error corrected and identified from fastq files, and data was aligned to the hg19 reference genome using CellRanger V2.1 Command Line tools. For measuring expression of neo selectable marker a second round of alignment was conducted to the hg19 reference genome along with the neomycin sequence from pLNCX2. Secondary analysis was conducted using Seruat R package version 3 [21, 22]. Data generated from each cell population (Growing, RS, OIS, and DDIS cells) was filtered seperatley. Stringent filtering methods were applied using parameters described in literature such as number of genes detected, number of unique transcripts detected, percent mitochondrial genes detected, and percent ribosomal RNA detected [21, 22]. To filter cells we refrained from regressing out the effects of the above-mentioned QC metrics when implementing clustering protocols. Then, we were able to cluster the majority of ‘low quality’ cells seperatley from the good quality cells. Low quality clusters were then discarded.

### Clustering scRNA-seq data

Filtered data sets were merged together, scaled, and re-normalized using Seurat [21, 22]. In this case the number of genes detected, number of unique transcripts detected, and percent of mitochondrial genes detected were all scaled out as described by the Seurat protocol, thereby ensuring that any conclusions drawn from downstream analysis were driven by variable gene expression and not technical factors such as cell to cell variability in sequencing depth. Next, principal component analysis (PCA) was conducted, and, depending on the data set, the top 20 to 30 principal components were fed into a SNN clustering algorithm.

### Differential Expression Analysis

Differential expression was performed using Seurat with MAST methodology [23, 24]. Cluster-1 was compared to Cluster- 2 in each type of senescence.

### Ingenuity Pathway Analysis

The computed log fold changes of significantly differentially expressed genes (false discovery rate less than 0.05) were used in Ingenuity Pathway Analysis (IPA) by Qiagen. Upstream regulators were plotted according to their z-score and q-value of their overlap using GGplot2 R package. Q-values for upstream regulators were computed using p.adjust function in base R.

### Gene Set Enrichment Analysis

Computed log fold changes of genes differentially expressed between Cluster-1 and Cluster-2 were used in a pre-ranked GSEA analysis. GO terms and KEGG pathways were downloaded from MSigDB. GO terms included custom SASP lists were based on existing literature [28–34]. KEGG pathways for primary and secondary senescence were generated by Teo et al. and were added to the list of KEGG pathways [11]. Bar charts were generated using ggplot2.

### Navitoclax Experiments and Analysis

Senescent cells were treated with 1 uM of Navitoclax for 3 days. For OIS and DDIS samples, 4OHT and Etoposide were removed from the growth media before Navitoclax treatment. DMSO treated cells served as controls. Cells recovered in regular growth media supplemented with DMSO overnight before being harvested for scRNA-seq. Clusters were identified in DMSO treated controls. P values were calculated using the chisq.test function in R.

### Retroviral Infections of ERT2-HRAS:G12V

293T cells were co-transfected with plasmid DNAs of a retroviral vector and the helper vectors using FuGENE HD (Promega). Medium was collected 24, 36, and 48 hr later for infection of LF1 cells. The vector backbone was clonetech pQCXIN (Addgene Cat Number 631514). Clonal cell lines were generated through serial dilution in 500 micrograms/ml of G418. Colonies generated from single cells were selected and further propagated.

### Cell Culture and Senescence Induction

We sequenced proliferating female human diploid fibroblast cells (LF1 cells) along with populations of LF1 cells that were induced into replicative senescence, oncogene induced senescence, and DNA damaged induced senescence. LF1 cells were obtained from the Sedivy lab. All cell populations were generated with a clonal cell line that possessed an ERT2- hRAS:G12V transgene. For proliferating populations cells were passaged in regular growth media and harvested at 60% to 80% confluency. Meanwhile replicative senescent cells were passaged in regular growth media until replicative exhaustion. DDIS was induced with addition of etoposide at 40uM for 3 weeks to regular growth media. OIS was induced by adding 4-Hydroxytamoxifen to regular growth media, at which point the cells underwent a hyperproliferative phase before senescing after 6 days. Regular Growth media consisted of HAMS F10, 15% FBS, and 1x of Penicillin Streptomycin and Glutamine. Cells were maintained at 37 degrees Celsius at 5% CO_2_ and 2.5 % O_2_.

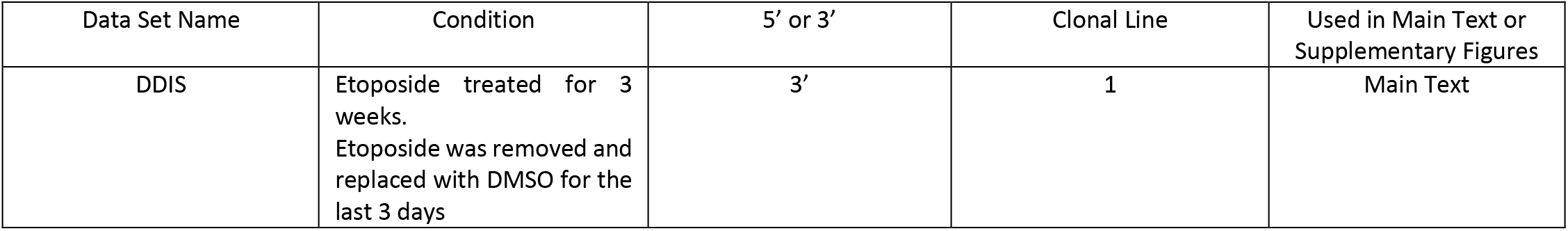
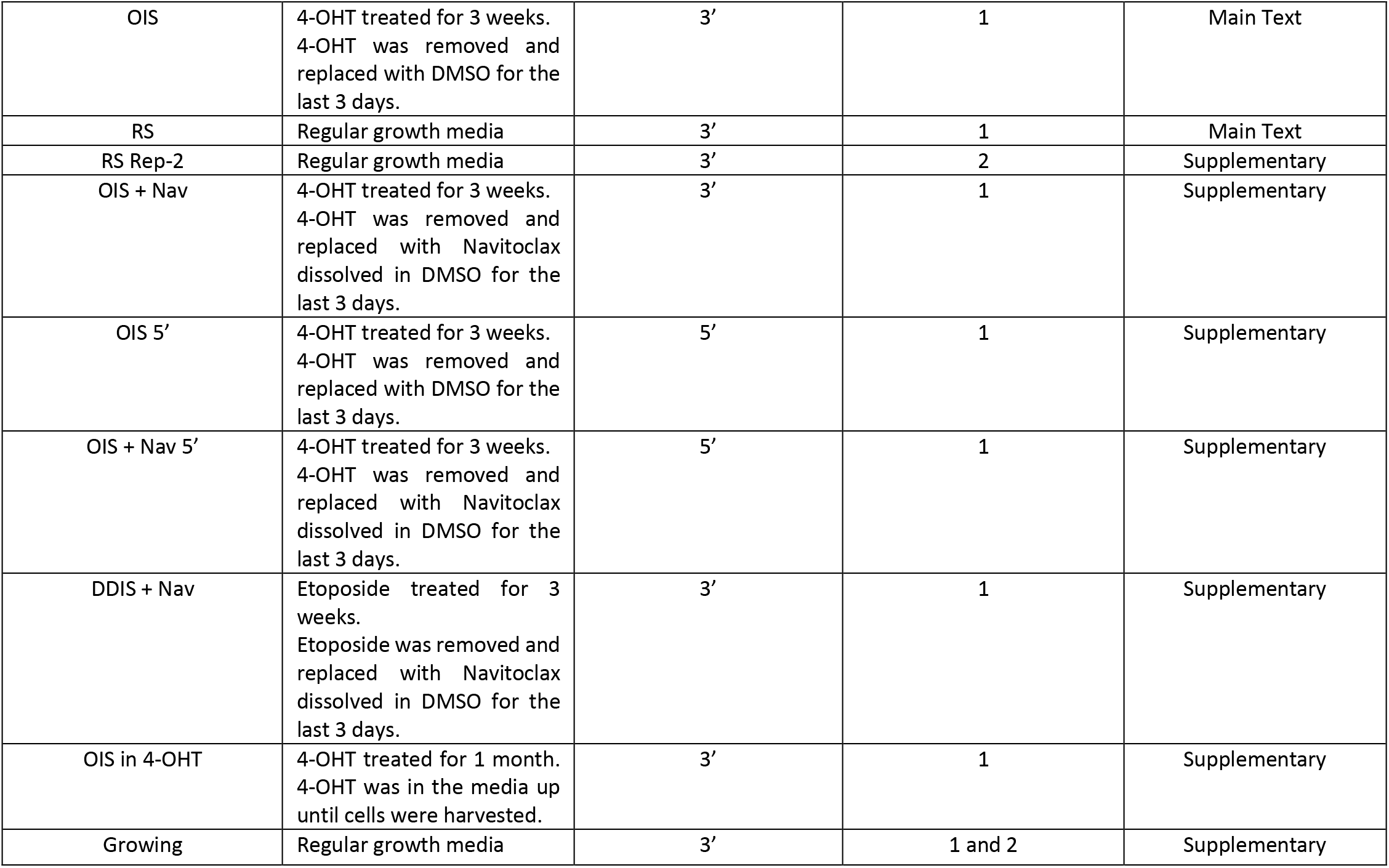

| Data Set Name | Condition | 5' or 3' | Clonal Line | Used in Main Text or Supplementary Figures |
| --- | --- | --- | --- | --- |
| DDIS | Etoposide treated for 3 weeks.<br>Etoposide was removed and replaced with DMSO for the last 3 days | 3' | 1 | Main Text |
| OIS | 4-OHT treated for 3 weeks.<br>4-OHT was removed and replaced with DMSO for the last 3 days. | 3' | 1 | Main Text |
| RS | Regular growth media | 3' | 1 | Main Text |
| RS Rep-2 | Regular growth media | 3' | 2 | Supplementary |
| OIS + Nav | 4-OHT treated for 3 weeks.<br>4-OHT was removed and replaced with Navitoclax dissolved in DMSO for the last 3 days. | 3' | 1 | Supplementary |
| OIS 5' | 4-OHT treated for 3 weeks.<br>4-OHT was removed and replaced with DMSO for the last 3 days. | 5' | 1 | Supplementary |
| OIS + Nav 5' | 4-OHT treated for 3 weeks.<br>4-OHT was removed and replaced with Navitoclax dissolved in DMSO for the last 3 days. | 5' | 1 | Supplementary |
| DDIS + Nav | Etoposide treated for 3 weeks.<br>Etoposide was removed and replaced with Navitoclax dissolved in DMSO for the last 3 days. | 3' | 1 | Supplementary |
| OIS in 4-OHT | 4-OHT treated for 1 month.<br>4-OHT was in the media up until cells were harvested. | 3' | 1 | Supplementary |
| Growing | Regular growth media | 3' | 1 and 2 | Supplementary |

## Acknowledgements

The N.N. lab was supported by IDeA grant P20GM109035 (Center for Computational Biology of Human Disease) from NIH NIGMS and grant 1R01AG050582-01A1 from NIH NIA. The content is solely the responsibility of the authors and does not necessarily represent the official views of the National Institutes of Health or any other organization. Figure models were created with BioRender.com.

## Conflict of interest

The authors declare no competing conflict of interests.

---

**Supplemental Figure 1.**
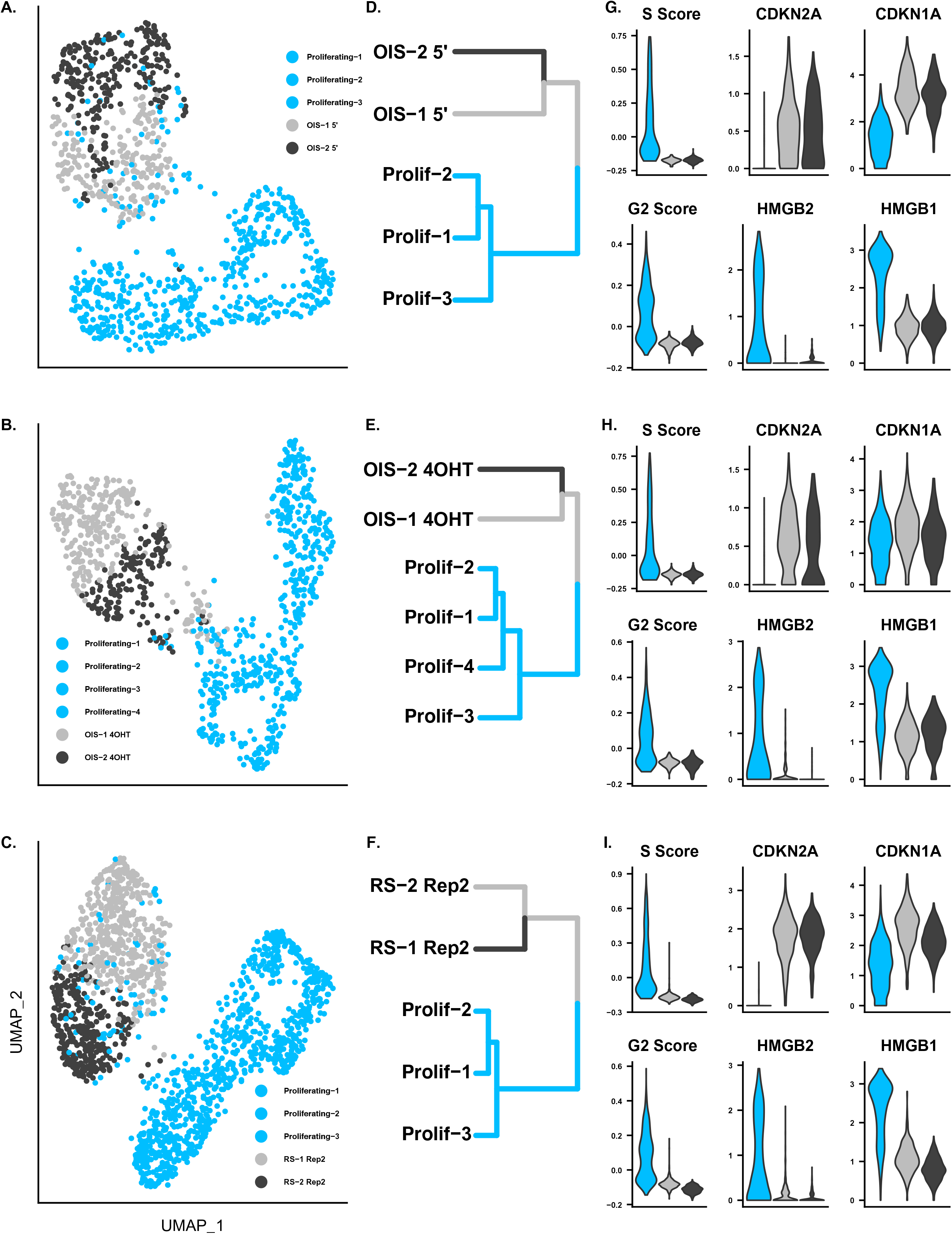
Clustering for Remaining and Senescent Markers for Cells Not Treated with Navitoclax A. - C. UMAP plots for all other data sets that were not treated with Navitoclax including OIS cells that were taken off of 4-OHT before collection and were sequenced from the 5’ end, OIS cells that were in 4-OHT when they were collected, and RS cells that were generated from a different clonal cell line than the rest of the data sets. **D. - F.** Cluster trees for the data sets. **G. - I.** Senescence markers for cluster-1 and cluster-2 for each data set.

**Supplemental Figure 2.**
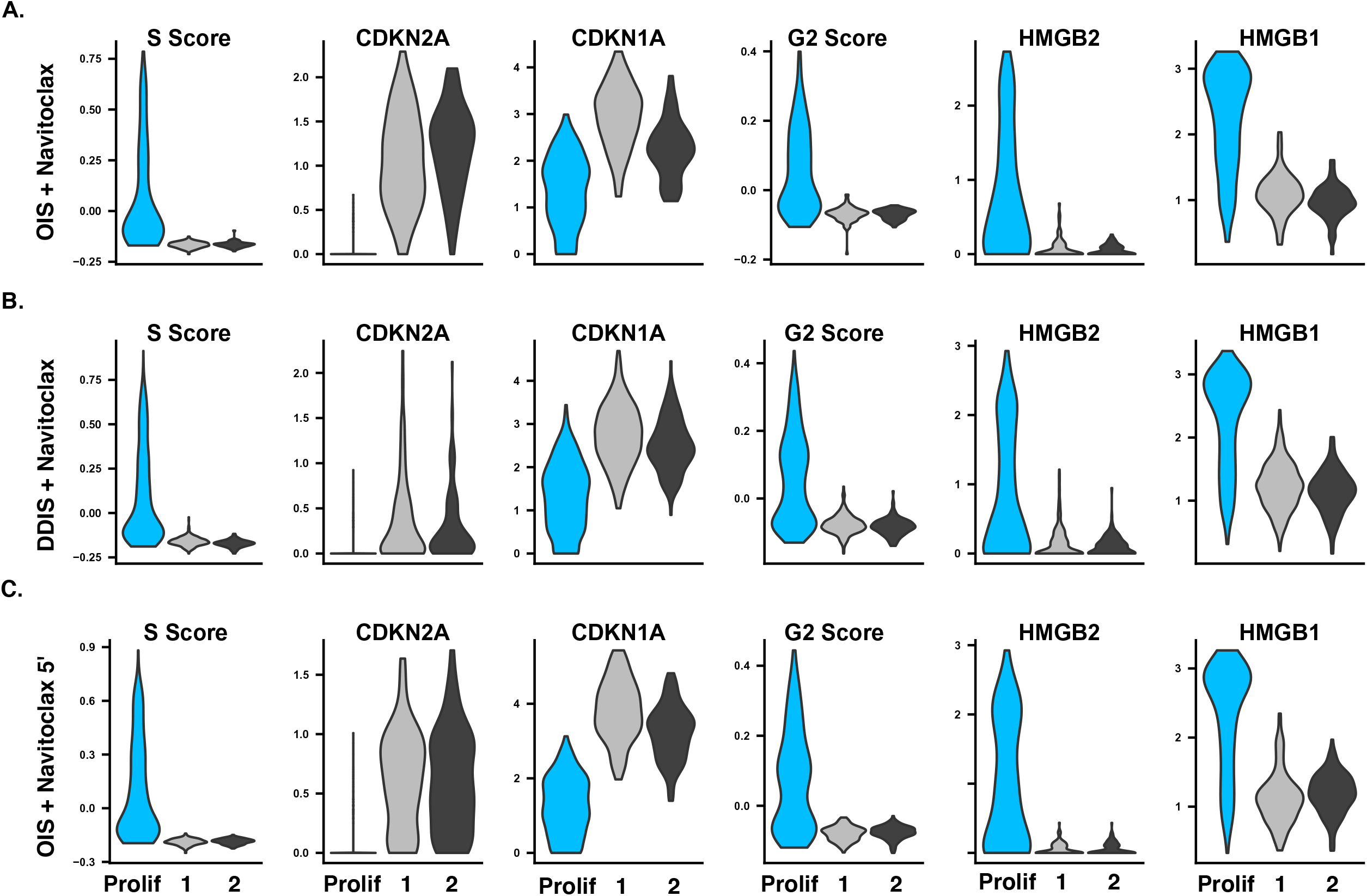
Senescent Markers for Navitoclax Treated Cells. **A. - C.** Senescence markers for cluster -1 and cluster -2 for all Navitoclax treated samples.

**Supplemental Figure 3.**
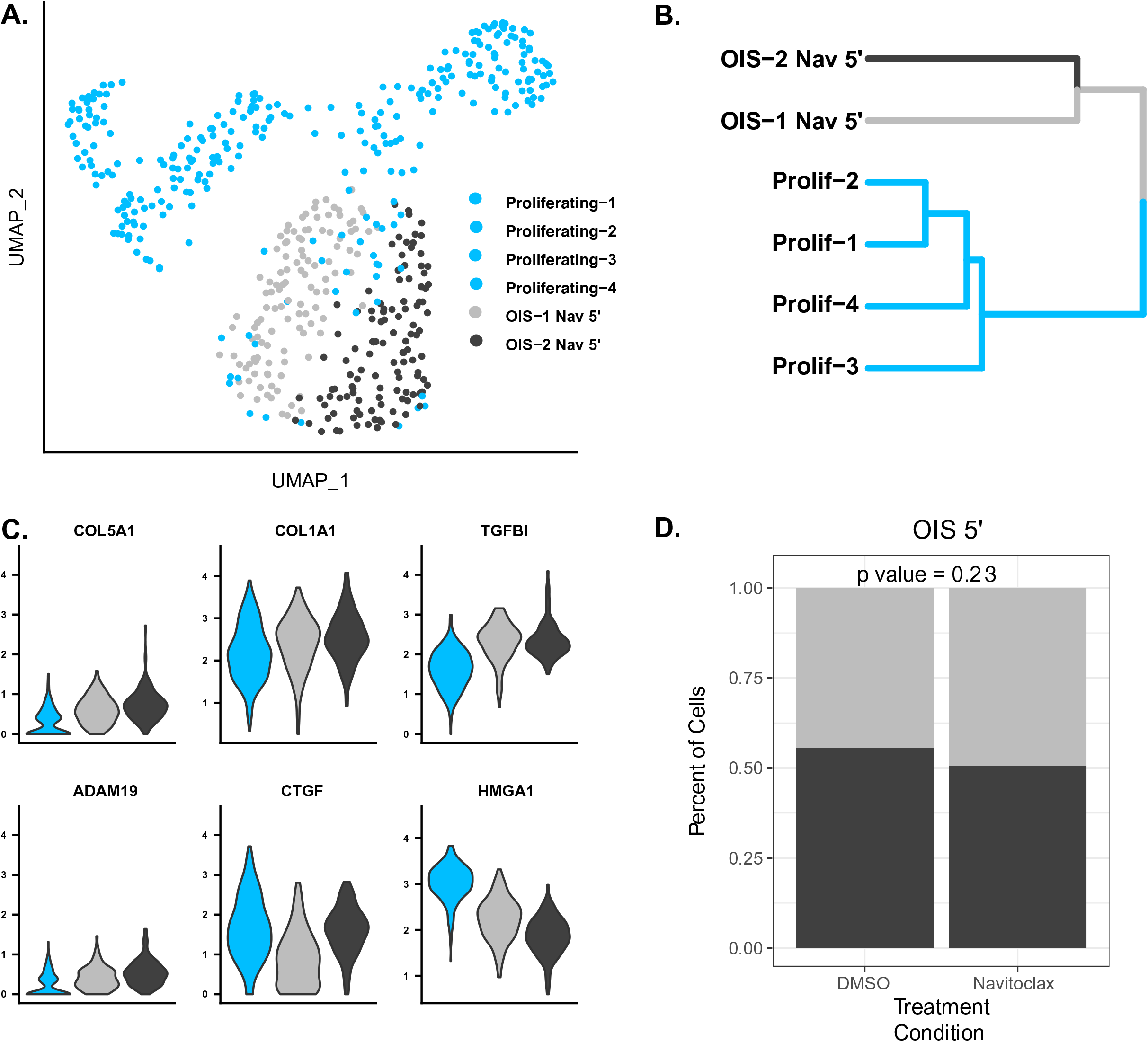
Navitoclax Preferentially Kills Cells in Cluster-1 in OIS Cells (Sequenced from 5’ End) **A.** UMAP plots for OIS cells treated with Navitoclax that were sequenced from the 5’ end. **B.** Cluster tree for that data set. **C.** Violin plots showing marker genes for primary and secondary senescence. **D.** Bar plots showing Navitoclax preferrtially induces apoptosis in Cluster-1 cells.

**Supplemental Figure 4.**
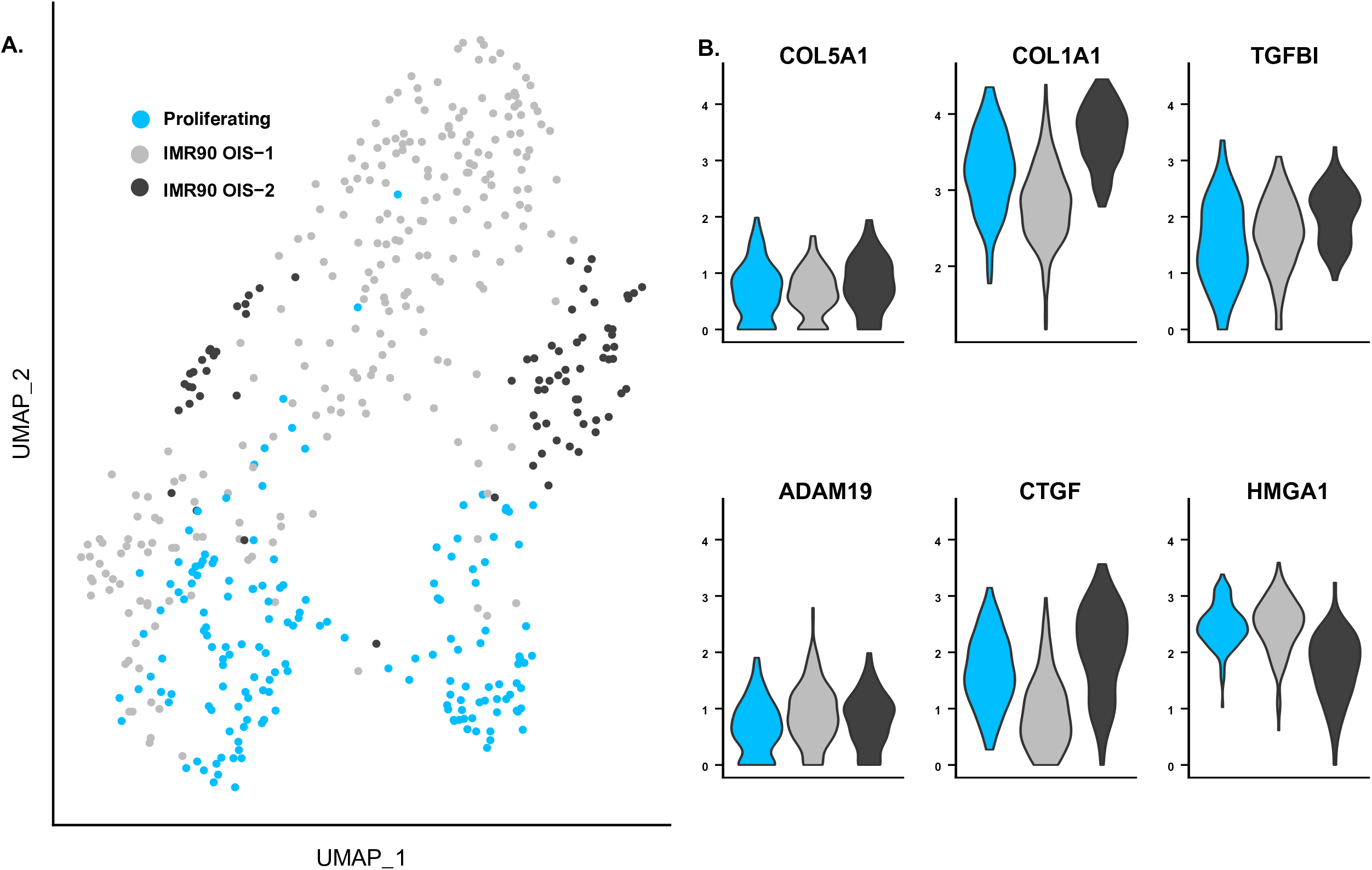
Primary and Secondary Markers From Non-Clonal OIS Cells. **A.** UMAP plots showing primary and secondary clusters for IMR90 cells from Teo et al. **B.** Primary and secondary marker genes for Teo et al.

**Supplemental Figure 5.**
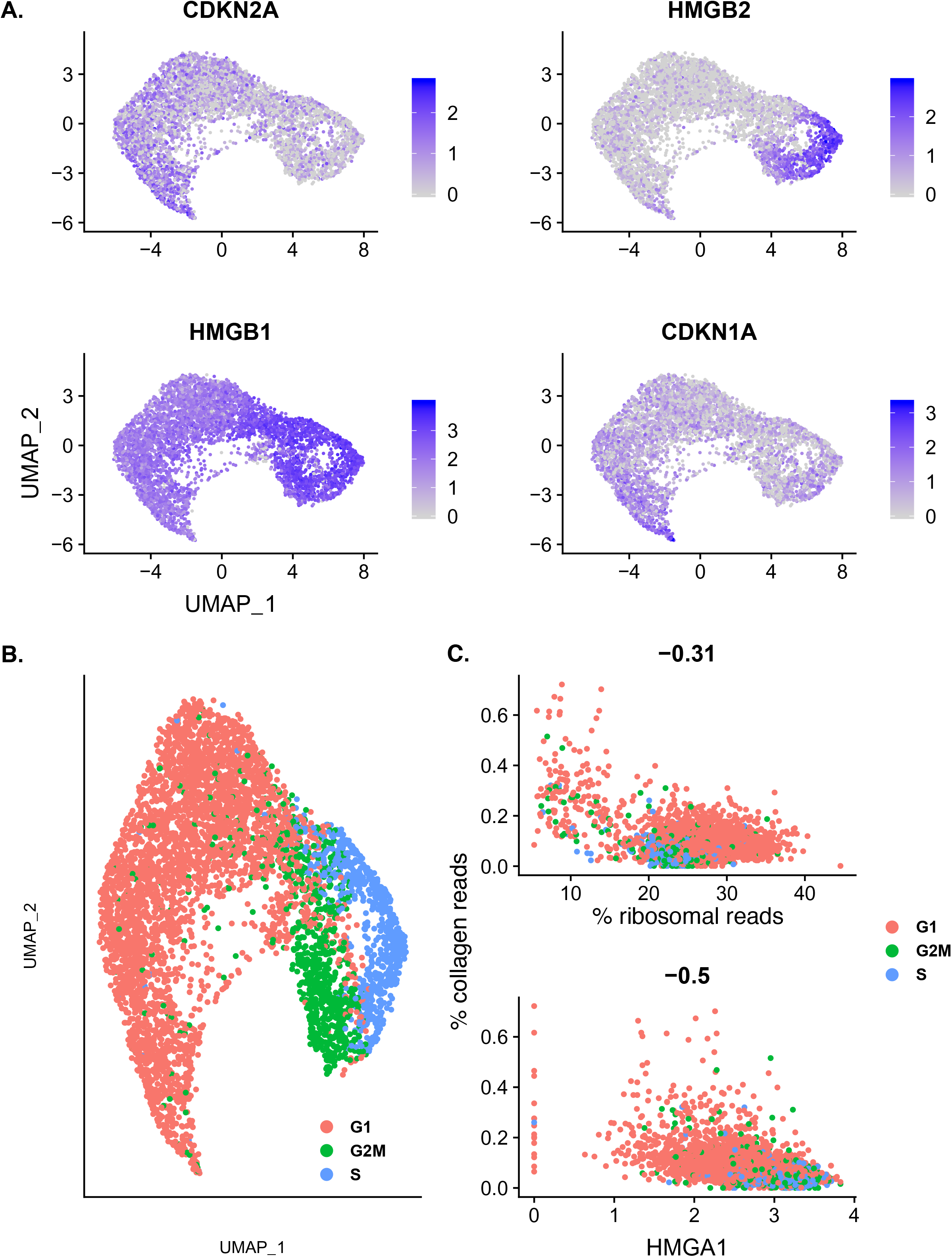
Cluster-1 and Cluster-2 Expression Profiles are Also Present in HUVEC RS Cells. Plots for HUVEC RS cells. **A.** Feature plots showing the expression of senescence markers in the data set. **B.** Predicted cell cycle phase. **C.** (Top) Scatter plot with cells plotted according to their ribosomal RNA expression on the x-axis and collagen expression on the y-axis. (Bottom) Scatter plot with cells plotted according to their HMGA1 expression on the x-axis and collagen expression on the y-axis.

**Supplemental Figure 6.**
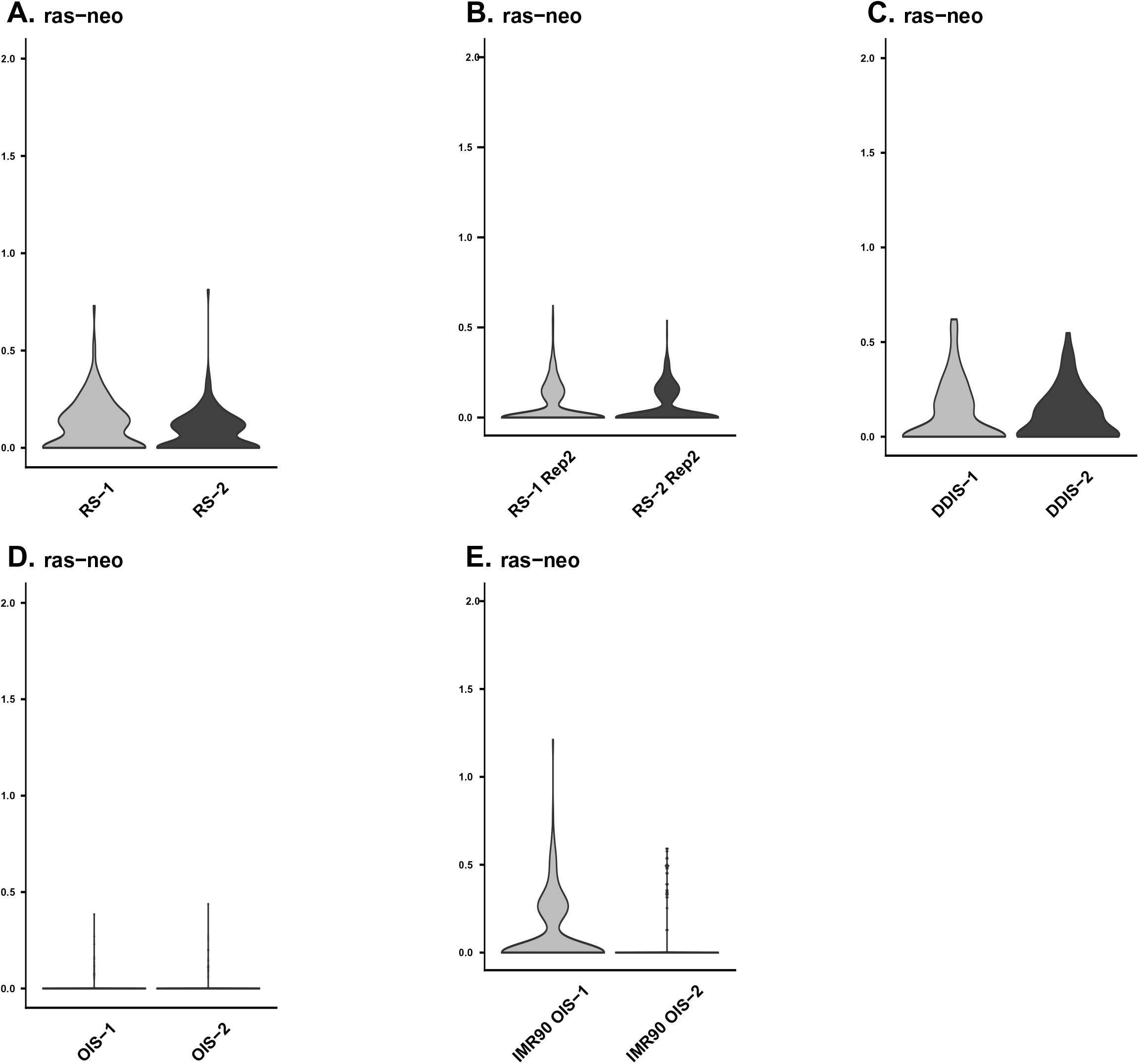
Clonal Cell Lines Express Similar Levels of 4-OHT inducible HRAS:G12V Transgene Between Subpopulations. **A. - C.** Violin plots showing expression of the neomycin resistance and 4-OHT inducible HRAS:G12V transgene between cluster-1 and cluster-2 for RS, OIS, and DDIS cells. **D.** Violin plots showing expression of the neomycin resistance and 4- OHT inducible HRAS:G12V transgene between cluster-1 and cluster-2 for IMR90 OIS cells from the Teo et al study.

**Supplemental Figure 7.**
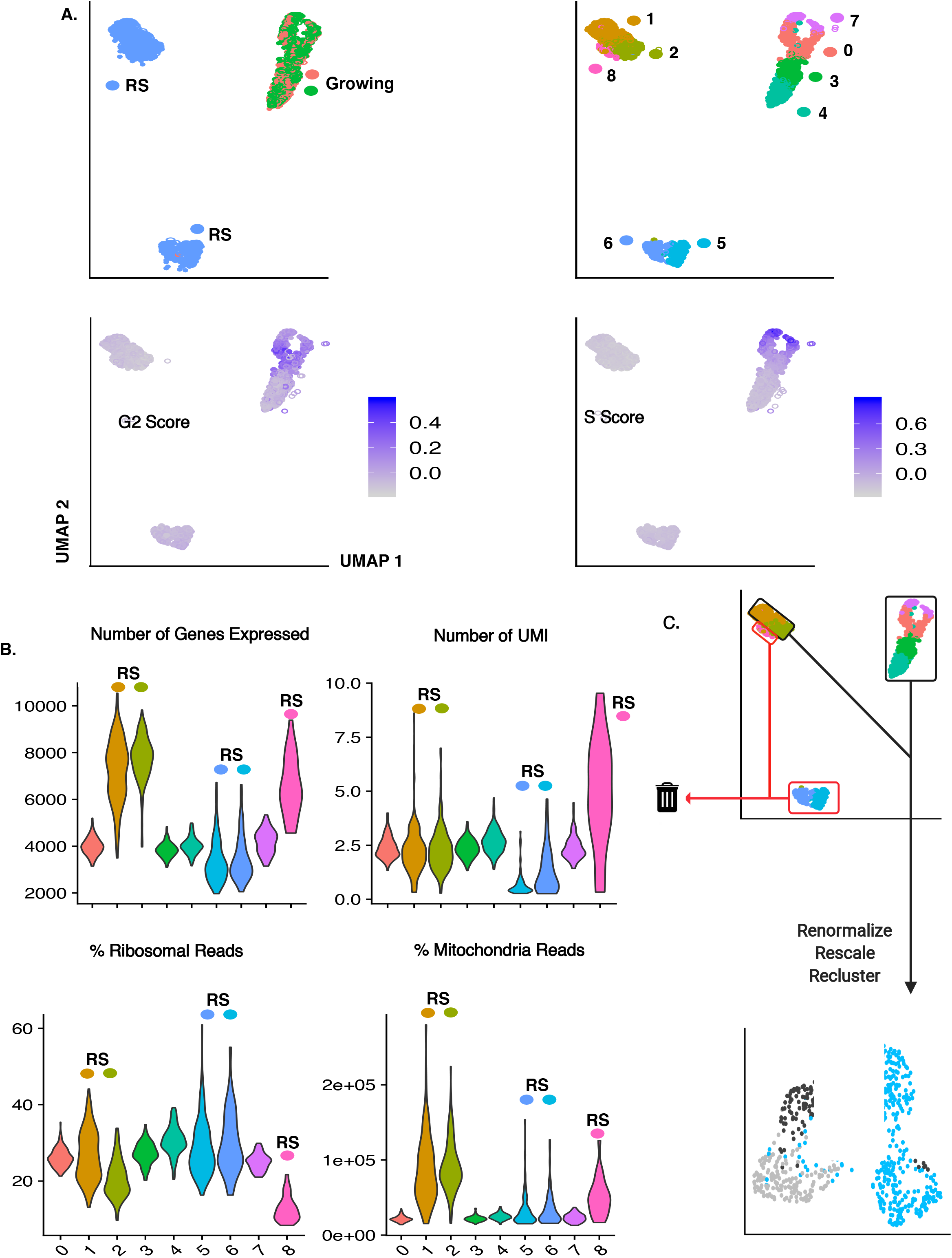
Filtering Method Applied To Data Sets. **A.** UMAP plots for RS cells generated from the second clonal line (RS Rep2). Top left shows samples. Top right shows identified clusters. Bottom left shows cell cycle score for the G2 phase of the cell cycle. Bottom right shows the cell cycle score for the S phase of the cell cycle. **B.** Violin plots showing QC parameters for the identified clusters. **C.** Schematic showing how filtering was performed.

**Supplemental Figure 8.**
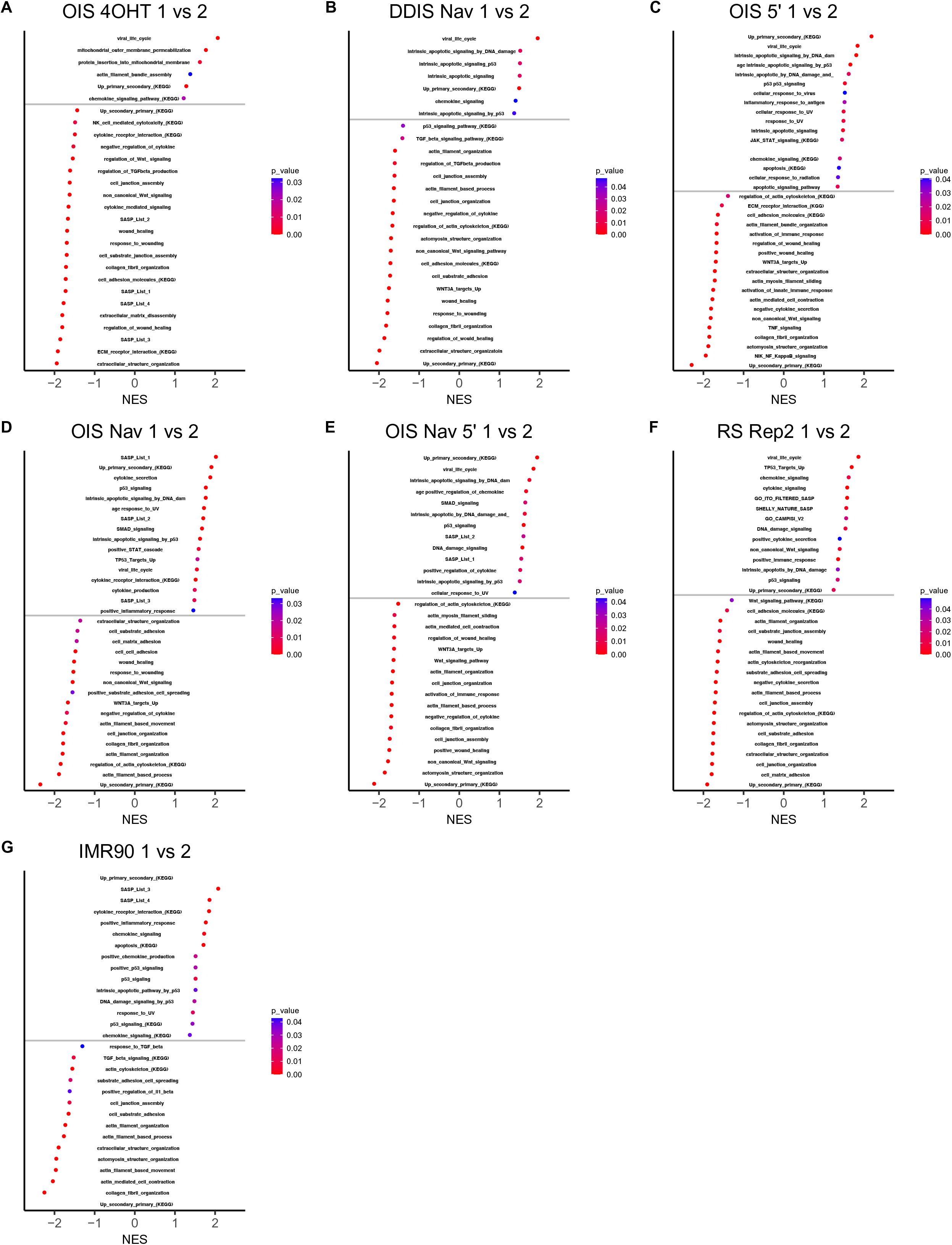
GSEA and KEGG Terms for all Other Data Sets and IMR90 OIS Cells from Teo et al. GSEA and KEGG plots comparing cluster -1 and cluster-2 for the remaining data sets in this study, in addition to those for the IMR90 OIS experiments comparing primary and secondary cells from Teo et al.

**Supplemental Figure 9.**
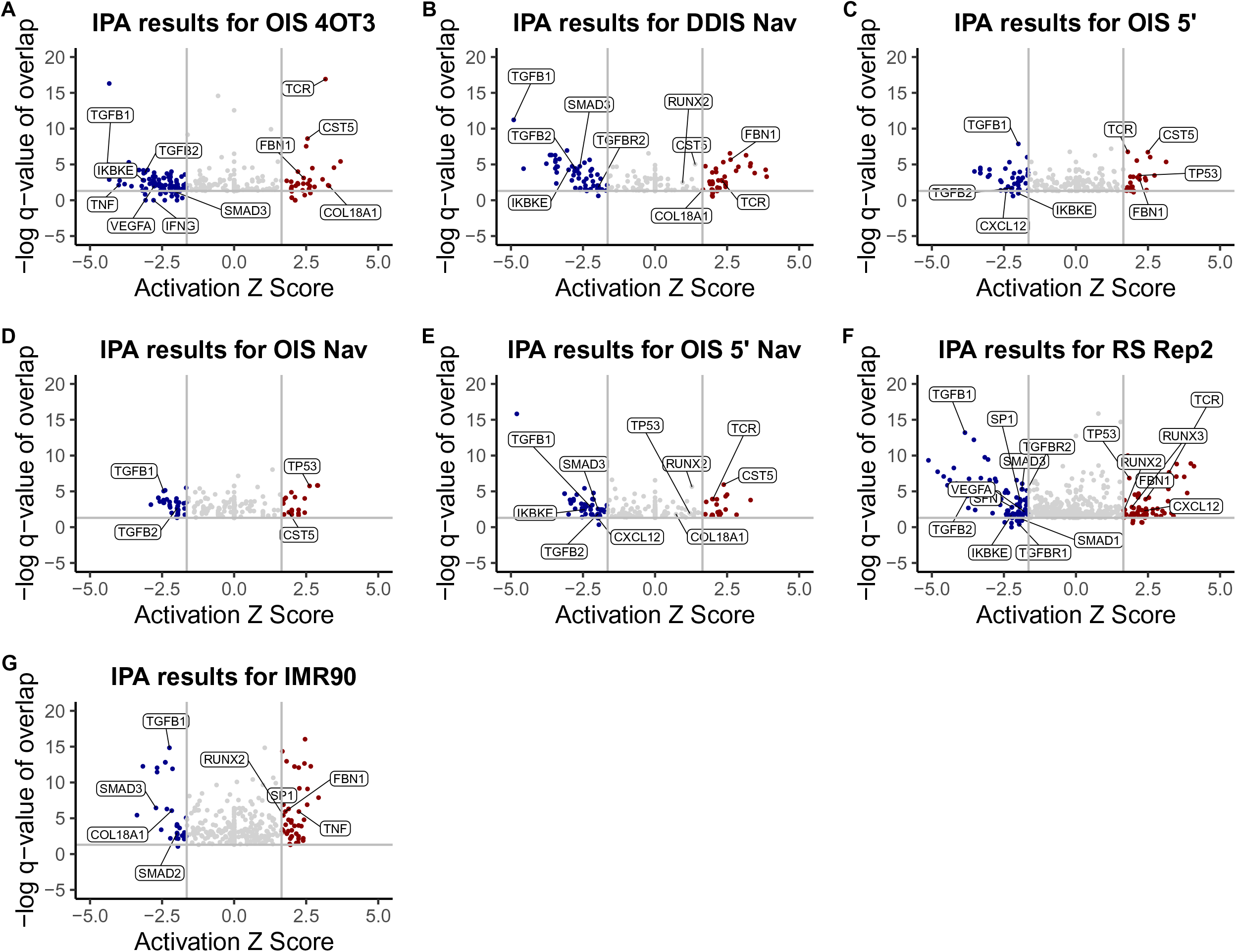
GSEA and KEGG Terms for all Other Data Sets and IMR90 OIS Cells from Teo et al. Voclano pots showing predicted upstream regulators (IPA) for cluster-1 (red) and cluster-2 (blue) for each type of senescence. Regulators are plotted according to their z-score on the x-axis which shows if they regulate cluster-1 or cluster-2. The ‘-log of the q-value of overlap’ is plotted on the y-axis. This is the negative logarithmic transformation of their false discovery rate, which determines the upstream regulators whose gene sets show a statistically significant overlap with the list of differentially expressed genes in the data. Plots for IMR90 cells from Teo et al are also shown.

## Notes

### Competing Interest Statement

The authors have declared no competing interest.

